# Shear-sensitive adhesion enables size-independent adhesive performance in stick insects

**DOI:** 10.1101/724831

**Authors:** David Labonte, Marie-Yon Struecker, Aleksandra Birn-Jeffery, Walter Federle

**Affiliations:** Department of Bioengineering, Imperial College, United Kingdom; Department of Zoology, University of Cambridge, United Kingdom; School of Engineering and Materials Science, Queen Mary University of London, United Kingdom

**Author notes:** Requests for further information and resources should be directed to and will be fulfilled by the Lead Contact, David Labonte.

## Abstract

The ability to climb with adhesive pads conveys significant advantages, and is hence widespread in the animal kingdom. The physics of adhesion predict that attachment is more challenging for large animals, whereas detachment is harder for small animals, due to the difference in surface-to-volume ratios. Here, we use stick insects to show that this problem is solved at both ends of the scale by linking adhesion to the applied shear force. Adhesive forces of individual insect pads, measured with perpendicular pull-offs, increased approximately in proportion to a linear pad dimension across instars. In sharp contrast, whole-body force measurements suggested area-scaling of adhesion. This discrepancy is explained by the presence of shear forces during whole-body measurements, as confirmed in experiments with pads sheared prior to detachment. When we applied shear forces proportional to either pad area or body weight, pad adhesion also scaled approximately with area or mass, respectively, providing a mechanism that can compensate for the size-related loss of adhesive performance predicted by isometry. We demonstrate that the adhesion-enhancing effect of shear forces is linked to pad sliding, which increased the maximum adhesive force per area sustainable by the pads. As shear forces in natural conditions are expected to scale with mass, sliding is more frequent and extensive in large animals, thus ensuring that large animals can attach safely, while small animals can still detach their pads effortlessly. Our results therefore help to explain how nature’s climbers maintain a dynamic attachment performance across seven orders of magnitude in body weight.

## Introduction

Many arthropods and small vertebrates use adhesive pads for climbing. These animals cover approximately seven orders of magnitude in body weight. Safe and effective climbing across such enormous size differences requires that the employed adhesive systems maintain performance even over large areas of contact, which is a fundamental challenge in adhesion science. In technical adhesives, this challenge typically arises from stress concentrations, which cause adhesive force per area (mean adhesive stress) to decrease as contact area increases, *σ* ∝ *A*^*≤*0^ [1, 2]. As an illustrative example, the force required to peel a thin strip of adhesive tape does not change whether the tape has a length of 2 cm or 500 km; instead, the peel force is proportional to a characteristic length – in this case the width of the tape – as all normal stresses are concentrated at the peeling edge [3]. In biological adhesive systems, this problem is exacerbated further, because the mass which needs to be supported is proportional to a volume, and hence scales as the cube of a characteristic length, *m ∝ V ∝ L*^3^. Adhesive pad area, in turn, is expected to grow more slowly, *A ∝ L*^2^ *∝ m*^2*/*3^, assuming geometric similarity (or ‘isometry’). Emerging from this simple geometrical argument is thus a non-trivial problem: Size-independent adhesive performance requires that the ratio of the maximum adhesive force an animal can sustain to its body mass, *m*, is constant:

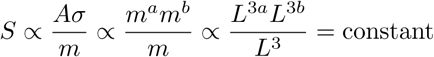

Here, *S* is a ‘safety factor’, and *a, b* are the scaling coefficients describing the relationship between pad area, adhesive stress and body mass, respectively. Size-independent performance requires *a* + *b* = 1, so that larger animals need to either (i) systemically increase the adhesive stress their pads can sustain (*b >* 0; ‘efficiency’); (ii) break with the condition of geometric similarity, and systematically increase the fraction of body surface area used for adhesive pads (*a >* 2*/*3; ‘positive allometry’); or (iii) combine both strategies [2]. What do climbing animals do?

Perhaps surprisingly, it appears that climbing animals make use of both strategies, albeit at different phylogenetic levels: Across distantly related animals, pad area grows in direct proportion to mass, *A ∝ L*^3^ *∝ m*, whereas it is approximately isometric within more closely related groups, *A ∝ l*^2^ *∝ m*^2*/*3^ [4]. This strong pylogenetic signal presumably reflects phylogenetic and developmental constraints: the fraction of the available surface area that is used for adhesive pads differs by a factor of about 200 between large geckos and tiny mites, and such extreme differences require substantial anatomical changes, which in turn may only be possible over long evolutionary time scales.

While strong positive allometry hence provides a partial answer to the puzzle of how climbing animals maintain adhesive performance, it leaves unresolved the question of whether and if so how animals within closely related groups compensate for the decrease in safety factor predicted by the isometric growth of their adhesive pads. Strikingly, there is robust evidence that within closely related groups, pad efficiency increases, and this increase can indeed be large enough to achieve constant safety factors despite an isometric growth of pad area, i. e. *σ ∝ L ∝ m*^1*/*3^ [2, 4, 5]. However, the mechanisms underlying this biologically important increase in pad efficiency have remained entirely unclear [though several authors have suggested corresponding hypotheses 2, 4–7].

In this article, we show that an increase in pad efficiency can arise as a direct consequence of the coupling between adhesive and shear forces widespread in animal adhesive pads [2, 8–11], thereby providing new insights into how both small and large animals can climb effectively with sticky feet.

## Results and discussion

### Shear forces control scaling of adhesion

We used a centrifuge to measure whole-body adhesion performance on smooth glass across all instars of Indian stick insects [*Carausius morosus*, Sinety, 1901. See Fig. 1 A & B. For details on this method, see ref. 12]. Across more than two orders of magnitude in body mass, *m*, adhesive force scaled as *m*^0.69^ (95% CI (0.59 — 0.79), n=45), suggesting a direct proportionality to the *area* of the adhesive pads which is approximately isometric (see Fig. 2 A and Tab. 1 for statistics on pad allometry. All slopes cited in this study were obtained with ordinary least-squares regression, but the main conclusions are independent of the regression technique used, see Tab. S1). However, the scaling coefficient of adhesion changed dramatically when forces of individual pads were measured by performing perpendicular pull-offs [Fig. 1 C. See ref. 13]. In sharp contrast to whole body measurements, single pad adhesive force scaled with *m*^0.34^ (95% CI (0.27 — 0.40), n=72), suggesting that it is proportional to a characteristic *length* of the isometric adhesive pads (see Fig 2 A). What is the origin of this discrepancy between single-pad and whole-body measurements?

**Table 1:** Scaling coefficients were obtained with both ordinary least squares and major axis regression. The effect of shear forces on the scaling of adhesive forces is consistent across both regression models, although the exact scaling coefficients differ slightly.

| *Ordinary least squares regression* | Slope | Elevation | Experimental condition |
| --- | --- | --- | --- |
| Single pad adhesion against mass | 0.34 (0.27, 0.40) | -1.63 (-1.77, -1.49) | No shear force |
| Single pad adhesion against mass | 0.71 (0.61, 0.82) | -1.91 (-2.13, -1.69) | Area-scaled shear force |
| Single pad adhesion against mass | 0.86 (0.70, 1.03) | -0.56 (-0.69, -0.42) | Mass-scaled shear force |
| Whole body adhesion against mass | 0.68 (0.61, 0.75) | -2.22 (-2.55, -1.90) | Shear force not controlled |
| Single pad adhesive stress against mass | 0.31 (0.15, 0.46) | 0.40 (0.11, 0.69) | Mass-scaled shear force |
| Contact area against mass | 0.68 (0.62, 0.74) | 3.12 (3.00, 3.24) | Area-scaled normal load |

| *Reduced major axis regression* | Slope | Elevation | Condition |
| --- | --- | --- | --- |
| Single pad adhesion against mass | 0.43 (0.37, 0.50) | -1.82 (-1.96, -1.68) | No shear force |
| Single pad adhesion against mass | 0.76 (0.7, 0.88) | -2.01 (-2.23, -1.80) | Area-scaled shear force |
| Single pad adhesion against mass | 0.94 (0.79, 1.23) | -0.62 (-0.76, -0.49) | Mass-scaled shear force |
| Whole body adhesion against mass | 0.71 (0.65, 0.79) | -2.36 (-2.69, -2.04) | Shear force not controlled |
| Single pad adhesive stress against mass | 0.45 (0.33, 0.62) | 0.13 (-0.16, 0.43) | Mass-scaled shear force |
| Contact area against mass | 0.70 (0.64, 0.76) | 3.08 (2.96, 3.20) | Area-scaled normal load |

**Figure 1:**
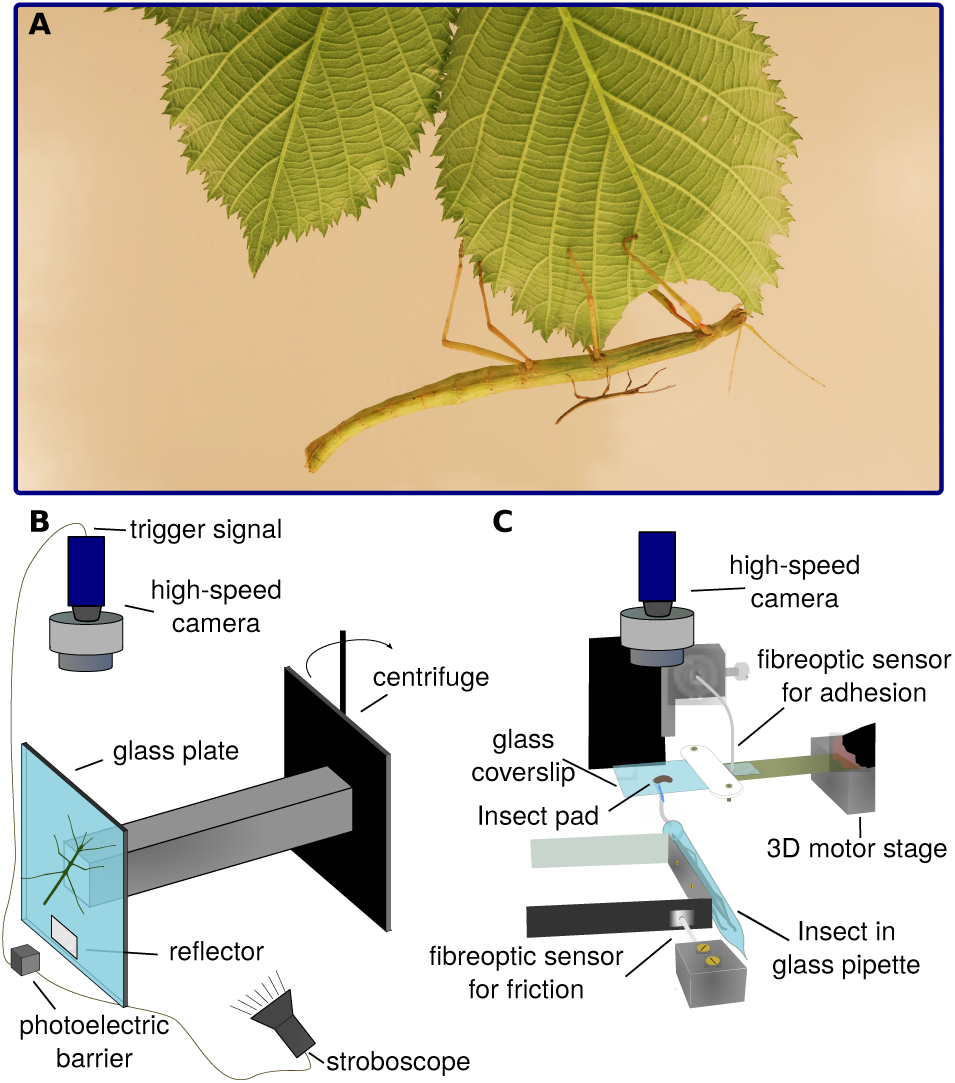
(A) Indian stick insects vary by almost three orders of magnitude in body mass (image S Chen). We measured the attachment performance across all instars using (B) a centrifuge, and (C) a custom-built 2D fibreoptic set-up.

**Figure 2:**
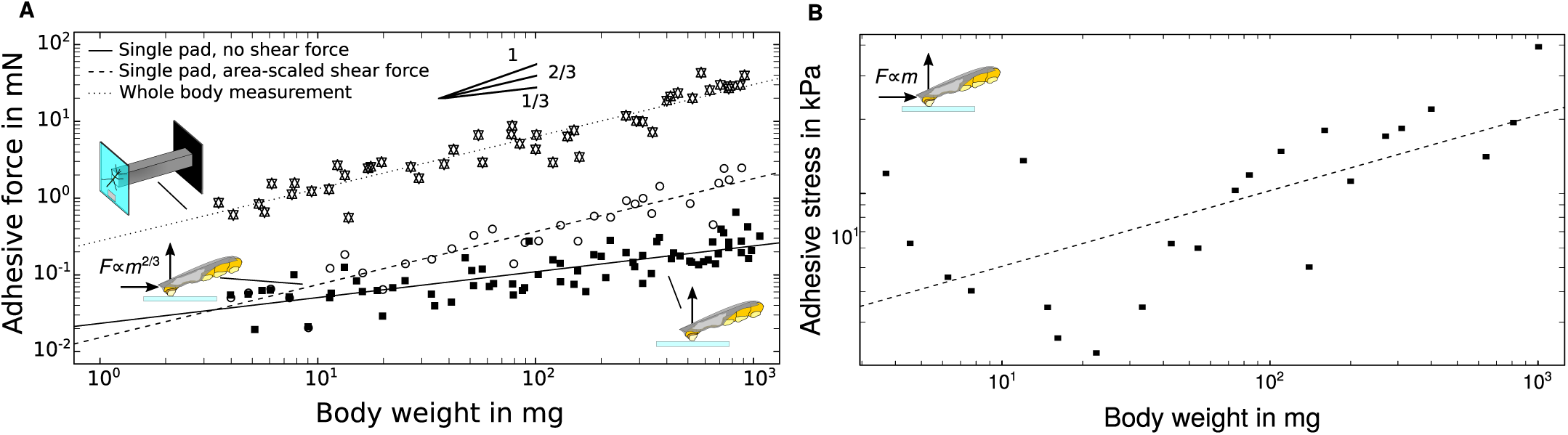
(A) Across all instars, whole-body adhesive performance scaled as *m*^0.69^ (95% CI (0.59 — 0.79), n=45; dotted line), whereas adhesive forces measured with perpendicular pull-offs of individual pads scaled as *m*^0.34^ (95% CI (0.27 — 0.40), n=72; solid line). The discrepancy arises from the absence of shear forces during perpendicular pull-offs. When shear forces scaled with pad area were applied prior to detachment, adhesive forces scaled as *m*^0.71^ (95% CI (0.61 — 0.82), n=32; dashed line). More detailed statistics can be found in the main text. (B) Because of the linear relationship between applied shear force and measured adhesion, applying a shear force corresponding to the insect’s body weight increases the scaling coefficient of adhesion, leading to an apparent increase in pad efficiency. When shear forces equivalent to one body weight were applied prior to detachment, adhesive stress increased with *m*^0.31^ (95% CI (0.16 — 0.45), n=23), sufficient to achieve size-independent safety factors. Note that both plots are double-logarithmic.

A key difference between the two measurements arises from the sprawled leg posture of stick insects, which results in an inward shear component of the force vector detaching individual pads during whole-body measurements. As a consequence, centrifuge measurements simultaneously induce both normal *and* shear stresses. Our single pad measurements, in contrast, only induced normal stresses. In order to investigate if the discrepancy between whole-animal and single-pad scaling arises from the presence or absence of shear forces, we repeated the single-pad measurements, but this time applied a feedback-controlled shear force to the pads prior to detachment [see ref. 14. for details on this method]. This shear force was scaled in proportion to *m*^0.67^ to achieve an approximately constant shear stress of about 25kPa, just above the static shear stress of the pads [*≈* 20 kPa, corresponding to a shear force of 0.1 mN for a stick insect of 5 mg weight, see ref. 10]. Single-pad adhesion forces measured in the presence of area-specific shear forces scaled as *m*^0.71^ (95% CI (0.61 — 0.82), n=32), virtually identical to the result obtained with whole-animal centrifuge measurements (ANCOVA, F1,73 = 0.36, p = 0.55, see Fig. 2 A), but significantly higher than for single-pad measurements in the absence of shear forces (ANCOVA, F1,100 = 40.81, p *<* 0.001, see Fig. 2 A). Hence, shear forces alter the scaling of adhesive forces, confirming our previous hypothesis [2].

Strikingly, our results also shed light on the conflict between the vast majority of theoretical adhesion models - which predict adhesive forces to grow more slowly than adhesive contact area [1, 2, 15] - and the majority of experimental data on biological adhesives - which imply area- or even above area-scaling [2]. This remarkable contradiction can be resolved by accounting for the size-related variation in shear forces acting during whole-body detachments, as we will show below. In the following, we will discuss why altering the magnitude and scaling of adhesion via shear forces is biologically important, and what mechanisms might explain the link between adhesion and shear force.

### Shear forces help to maintain

#### size-independent safety factors across body sizes

Animals climbing with adhesive pads vary by almost seven orders of magnitude in mass, which poses a significant challenge: the weight that needs to be supported grows faster than the area available for adhesive structures (assuming isometry). Across the entire size range of animals climbing with adhesive pads, this change in surface-to-volume ratio is predicted to reduce safety factors (i.e. adhesion per body weight) by a non-trivial factor of (10^7*/*3^) *≈* 200 [here, we also assumed area scaling of adhesion, and size-independent maximum adhesive stress. See 4]. Recently, we showed that heavier climbing animals partially solve this problem by allocating a larger fraction of the total available body surface area to adhesive structures [4]. However, this disproportionate increase in pad area did not occur within closely related taxa. Instead, mass-specific pad area differed considerably between vertebrates and invertebrates, but was approximately consistent with isometry within clades, indicating that pad size is constrained by phylogeny [4]. Strikingly, some vertebrate and invertebrate taxa are nevertheless able to achieve size-independent adhesion: their pads appear to get more effective as they grow in size [2, 4, 5]. Larger climbing animals hence appear to maintain size-independent safety factors by employing two distinct strategies – disproportionately larger versus more effective pads – at different phylogenetic levels [2, 4]. However, the underlying mechanism of the systematic increase in pad efficiency, observed within closely related groups [2, 4, 5], has remained unclear.

Based on the empirical observation that adhesive forces of invertebrate and vertebrate pads are an approximately linear function of the shear force that is acting during detachment [8, 10], one may speculate that shear forces can in principle be used to achieve an arbitrary scaling of adhesive forces, if they are varied systematically with size (with an upper limit set by the maximum sliding shear stress sustainable by the adhesive pads). In order to test this hypothesis, we conducted another series of single-pad measurements, this time applying a shear force equal to the weight of the individuals. The resulting adhesive force scaled with *m*^0.87^ (95% CI (0.70 — 1.03), n=23), an increase significantly exceeding that of adhesive pad area (*t*54= 2.5, p *<* 0.05). Force per pad area increased with *m*^0.31^ (95% CI (0.16 — 0.45), n=23, see Fig. 2 B), consistent with previous reports on tree frogs and ants [2, 4]. The relationship between shear force and adhesion may hence underlie the previously unexplained increases in pad efficiency with size [2, 4], and may even suffice to achieve size-independent safety factors (i.e. adhesion *∝ m* or adhesive stress *∝ m*^1*/*3^ for isometric animals). We therefore propose that the link between shear force and adhesion plays a key role not only for the controllability of attachment during locomotion [10, 16, 17], but also avoids the predicted decrease of safety factors in larger animals [2].

#### Biomechanics of shear-sensitive adhesion

While our data provide strong evidence that shear forces influence both the scaling *and* magnitude of adhesive forces [2], the physical basis for this effect remains unclear. Shearing pads towards the body is known to increase the adhesive contact area of smooth and hairy pads [18–20], but this does not explain the shear-dependence of adhesive stress observed here and in previous studies [14]. Numerous theoretical models have been proposed for the performance of animal adhesive pads [for a review, see for example 15], but to our knowledge no theory has been able to quantitatively predict the effect of shear forces on adhesion from first principles [see 10]. For example, the peeling theory for inextensible tape predicts that adhesion increases with applied shear force [see 2, 10, 15, 16, 21–23], and that it scales with length, seemingly consistent with our no-shear single-pad measurements (see Fig. 2 A). However, peeling theory fails to explain the well-established linear relationship between shear force and adhesion [10, 16], and is also inconsistent with the area-scaling of adhesive forces observed in pull-off measurements involving shear (see Fig. 2 A). This inconsistency may arise because shear forces increase the length of the peel zone, thereby leading to a more uniform stress distribution within the contact zone and hence to area scaling [2, 24, 25]. Notably, single-pad adhesion forces of stick insects follow peeling theory for small shear forces (or large peel angles), and only depart from theoretical predictions for large shear forces [or small peel angles, see 10]. The departure from peeling theory coincided with the onset of whole pad sliding during detachment [10], and two explanations for this observation have been proposed: First, pads that slide will be stretched, which increases their effective stiffness, thereby preventing the drop in adhesion predicted by the theory for extensible tape as peeling angles approach 0° [for a more detailed discussion, see 10, 23, 26]. While this effect is undoubtedly important, it cannot explain the departure from the theory for inextensible tape, which *underestimates* adhesion for small peel angles and still predicts adhesive forces to scale with a linear dimension of the contact [3, 10, 23]. Second, sliding results in the depletion of the contact-mediating liquid secreted by the adhesive pads [see Fig. 3 A. 10, 27, 28. For more details on the function and chemical composition of the secretion, see refs. [13, 29]]. Such a reduction in the amount of pad fluid has been hypothesised to increase pad adhesion, for example by increasing the contribution of ‘wet’ forces arising from the secretion’s surface tension and viscosity [19, 30], or by reducing the “interfacial mobility” [13]. However, direct evidence for a link between sliding, and changes in the force required to detach the pads is still missing.

**Figure 3:**
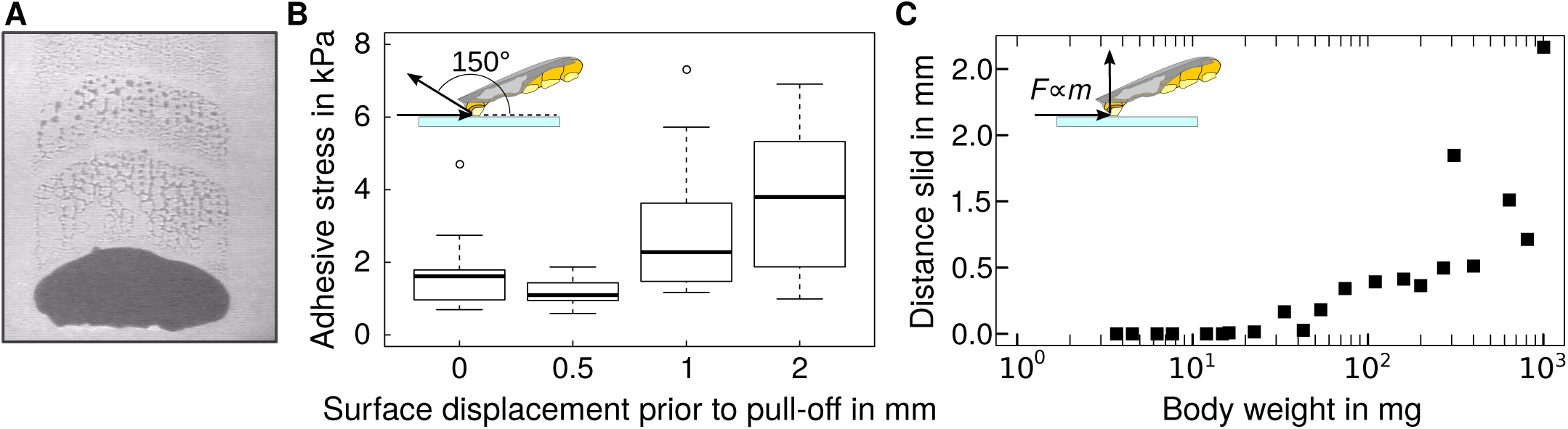
(A) Pads that slid left behind trails of contact-mediating fluid. (B) Adhesive force of pads that slid over different distances, prior to detachment at 150°, significantly increased with sliding distance (repeated measures ANCOVA, F1,37=20.1, p *<* 0.01, n = 10). (C) Sliding distance of stick insect pads before detachment in adhesion tests where shear forces equivalent to one body weight were applied. Because pad area grows less slowly than mass, larger insects are more likely to slide. Sliding occurred only for insects weighing more than approximately 20 mg, consistent with a simple estimate based on the pad allometry and the static shear stress (see main text). Because sliding increases adhesive strength, the variable sliding distance of pads helps large animals to attach safely, while small animals can still detach their pads effortlessly.

We performed a direct test of the hypothesis that pad sliding increases the strength of the adhesive contact. In brief, we conducted single-pad adhesion measurements in which pads of adult stick insects were initially subjected to proximal shear displacements of 0, 0.5, 1 or 2 mm, in a direction corresponding to a pull of the pad towards the body. Subsequently, the pads were detached with an angle of 150 ° relative to the surface. This experimental design allowed us to separate the effect of pad sliding from a possible effect of the shear force itself. Although the shear force acting on the pads at peak adhesion did not differ between treatments (repeated measures ANCOVA, F1,36=0.41, p = 0.53, n = 10), adhesion significantly increased with the applied shear displacement (repeated measures ANCOVA, F1,37=20.1, p *<* 0.01, n = 10), providing direct evidence for a sliding-induced change in interface strength [See Fig. 3 B. 10, 13].

#### The limits of shear-sensitive adhesion

We have demonstrated that shear forces control the magnitude of adhesive forces, and can therefore modify the scaling of adhesive forces. The adhesion-enhancing effect of shear force arises at least partly from pad sliding, which stretches the pad, and strengthens the contact. Does the amount of sliding differ between animals of different size, and if so, what are the consequences for the scaling of attachment performance?

Under natural conditions, pads will be sheared passively whenever adhesive forces are required, due to the insects’ sprawled leg posture. These passive shear forces likely scale with body mass, leading to a higher shear stress acting on the adhesive pads of larger animals, which are therefore more likely to slide. As an illustrative example, the shear stress acting on the pads of a 1000 mg adult stick insect in our single-pad experiments with mass-scaled shear forces was *≈* 70 kPa, or 3.5 times the static shear stress. For a 1^st^ instar insect weighing 5 mg, in turn, one body weight in shear force resulted in a stress of *≈* 10 kPa, about half the static shear stress. As a consequence, the transition to sliding occurred at some intermediate body mass, *mc*. We estimated *mc* using the allometric relationship of the adhesive pad area, and the approximate static shear stress of 20 kPA, yielding *mc* =21.8 mg. This estimate is in excellent agreement with direct observations of pad sliding extracted from video recordings (see Fig. 3 C). It is remarkable that the static shear stress of the pads is of the same order of magnitude as the stresses expected from one body weight acting on a single foot, as sliding is considered detrimental for the function of conventional adhesives (although under natural conditions, the stresses might be smaller as the force is shared between multiple pads). We believe this is no coincidence: the problem of surface attachment at varying body size can be seen from two perspectives: it is hard to detach when small, but challenging to attach when large. The magnitude of static shear stress of adhesive pads might hence be adaptive, as it can enable similar attachment performance for both small and large insects. Small animals are less likely to slide, and hence can more easily detach their pads. Large animals, in turn, benefit from the weight-specific adhesion enhancement caused by larger amounts of pad sliding, and hence are able to maintain adhesive performance. For pads tested at shear stresses in excess of their static shear stress, the distance slid increased as *m*^1.1^ (95% CI (0.75 — 1.43), n=15. See Fig. 3 C). At present, the quantitative relationship between the distance slid and the increase in adhesive strength is still unclear, and will be addressed in future work. It is plausible that adhesion enhancement depends on distance slid per unit pad length, adding further complexity to the scaling of shear-sensitive adhesion.

It appears relevant to distinguish between two types of sliding: (i) local sliding of parts of the pad’s contact zone near the peel front, which stretches the pad and thereby increases its stiffness, but does not lead to much fluid depletion; (ii) sliding of the *entire* pad, which will be most effective at depleting fluid by leaving it behind at the trailing edge [31]. This distinction can be used to illustrate a significant complication which we have thus far ignored for simplicity: During detachment, the contact area of the pads will decrease to zero, and the shear stress will therefore tend to notational infinity. Hence, even for a size-invariant shear stress that was initially below the static shear stress, the pads of all instars must slide at some point during detachment. If the shear stress exceeds the static shear stress only after peak adhesion has been reached, sliding will have no effect on adhesive performance. If pads slide prior to reaching the adhesion peak, however, the critical parameter governing attachment performance is the distance slid before detachment (Fig. 3 C). In insects, sliding speed increases in an approximately linear fashion with shear stress after the static shear stress is surpassed [32], but it is unclear if this relationship is affected by animal size. The time to detachment, in turn, depends on a number of factors, such as contact size and pad modulus, which ultimately together control the speed of crack propagation [13]. While a detailed investigation of the size-dependence of these factors will have to await future work, we point out that the need to generate some static shear stress provides a potential explanation for the increase in mass-specific pad area from invertebrate to vertebrate taxa [10]: From the previous discussion, it is unclear why vertebrates evolved pads much larger relative to their size, instead of just relying on the adhesion-enhancement provided by pad sliding. However, the maximum static shear stress sustainable by the pads is finite, and there hence must be a critical size at which the pad area must increase disproportionally to prevent large animals from sliding excessively. Future research should clarify the factors that determine sliding distance before detachment, its role for adhesion enhancement, as well as detachment dynamics across body sizes. Such work will ultimately further our understanding of how nature’s best climbers maintain performance across considerable variations in body size, and potentially allow us to transfer their tricks to scalable bio-inspired adhesives.

## Materials and Methods

### Experimental model organism

We used Indian stick insects (*Carausius morosus*) as model species, because they vary by almost three orders of magnitude in body weight between first-instar nymphs and adults Fig. 1 A. Individuals from all instars were taken from a laboratory colony kept at ambient conditions, and fed with water, ivy and bramble leaves *ad libitum*. All individuals were weighed to the nearest 10*µ*g (MC 5, Sartorius AG, Göttingen, Germany). Prior to further preparation, all distal pads (arolia) were investigated using light microscopy, to exclude individuals with damaged pads.

All experiments were conducted in ambient laboratory conditions (temperature: 19-23° C, relative humidity: 30-50%). In order to avoid pseudo-replication, all animals were kept in a different cage once tested.

Throughout this manuscript, we use ‘shear force’ to refer to forces applied parallel to the surface (they are - acted by ‘friction forces’). ‘Adhesive force’, in turn, refers to the normal component of the force resisting detachment for whole animals or individual adhesive pads. ‘Shear stress’ and ‘adhesive stress’ refer to these forces when normalised by contact area. We use ‘pad efficiency’ for the maximum adhesive stress a pad can produce. ‘Static shear stress’ is the maximum stress which can be applied parallel to the surface without causing the pads to slide, i. e. to move relative to the surface.

### Whole-body measurements

Whole-body adhesion measurements were conducted using a custom-built centrifuge [(Fig. 1 B) 12]. Animals were placed on vertical glass plates mounted on a custom-made holder attached to the centrifuge, which was gradually accelerated until the insects detached. A DMK 23UP1300 high-speed camera (Imaging Source Europe GmbH, Bremen, Germany) was mounted above the setup, and triggered by a photoelectric barrier to synchronise with the rotational speed of the centrifuge. To achieve sharp images, the setup was illuminated with a stroboscope, also triggered by the photoelectric barrier. We digitized the insect’s radial position on the centrifuge just before detachment, allowing us to calculate centrifugal acceleration; detachment force was then calculated as the product of body mass and centrifugal acceleration. Each insect was only measured once.

### Single-pad measurements

In order to isolate individual pads, stick insects were put into glass pipettes, and one of the two protruding front legs was attached onto a metal wire using dental wax (Elite HD, Zhermack, Badia Polesine, Italy), so that the ventral side of the arolium was facing up. To avoid interference with the measurements, the claw tips were cut off under a stereo-microscope (MZ16 Leica Camera AG, Wetzlar, Germany) using sharp tweezers, and dust particles were removed from the pad using a piece of sticky tape (Tesa SE, Norderstedt, Germany).

Single pad forces were measured with a custom-built 2D fibre-optic transducer set-up (Fig. 1 C). In order to eliminate cross-talk between the shear force and adhesion channels, they were physically separated: adhesion was measured by the deflection of a cantilever beam to which a glass coverslip was attached, and friction was measured by the deflection of an independent double cantilever beam to which a plastic tube holding the stick insect was attached (see Fig. 1 C). This separation also allowed a straightforward independent manipulation of the beams’ spring constants. For both beams, the deflection was sensed with fibre optic sensors (D12, Philtec, INC., Annapolis, USA), via small pieces of reflective foil glued to the far end of the beams (see Fig. 1 C). More details regarding the set-up and the calibration procedure can be found in [13].

In order to perform controlled experiments, the adhesion beam was mounted on a 3D motor positioning stage (M-126PD, Physik Instrumente, Karlsruhe, Germany), which was controlled with a custom-made Labview script [30]. A high-speed camera (DMK 23UP1300, The Imaging Source Europe GmbH, Bremen, Germany), mounted on top of a stereo-microscope (Wild M8, Wild Heerbrugg AG, Gais, Switzerland) allowed us to record the contact area of the pads during the measurements using reflected light. The output from both fibre-optic sensors, and the camera trigger signal were recorded at 1kHz with a data acquisition board (USB-6002, National Instruments, Austin, USA), allowing us to synchronise the contact area images with the measured forces.

For all single-pad measurements, pads were first brought in contact with clean glass coverslips for a period of 8 s, at a constant load of 0.5 mN unless otherwise specified [the normal load has no significant influence on adhesion measurements on smooth surfaces, see 14]. The normal load was kept constant using a force-feedback algorithm implemented in the LabView software [30], and the same algorithm was used to apply constant shear forces for a period of 10 s in all experiments involving controlled shear forces, detailed in the results section. All measurements ended with an upward movement of the motor stage holding the glass coverslip, at a speed of 0.5 mm s^*-*1^, which led to complete detachment of the pads.

### Quantification and statistical analysis

All force data were filtered with a low-pass Butterworth-filter in MatLabR2013a (Mathworks Inc., Natick, Massachusetts, USA), from which we extracted the peak adhesion force. The contact area and sliding distance of the pads were measured with ImageJv1.49 (National Institutes of Health, Bethesda, Maryland, USA). The video recordings were binarised by thresholding, and contact area was then extracted using native particle analysis routines. Scaling data was analysed using Ordinary least-Squares regression (OLS), because the error in the determination of mass is likely much smaller than that in the measurement of adhesive forces. There is some controversy as to whether OLS or standardised major axis (SMA) procedures are more appropriate for analysing scaling data [33–36], and we verified that all main conclusions of the paper hold independent of the regression technique, using the R-package smatr v3.4.4 [37]. The exact tests used, and sample size n, indicating number of individuals, are specified both in the results text and in the relevant figure captions. Effects were considered significant if p*<*0.05. Boxplots show the median and the 25%/75% quartiles; whiskers indicate 1.5— the interquartile range. All statistical analyses were performed in R v.3.4.4.

Requests for further information and resources should be directed to and will be fulfilled by the Lead Contact, David Labonte.

## Acknowledgements

This study was supported by research grants by the Biotechnology and Biological Sciences Research Council (BB/I008667/1 & B/R017360/1) to W.F. and D.L, respectively.

## References

1. Spolenak, R., Gorb, S., Gao, H., and Arzt, E. (2005). Effects of contact shape on the scaling of biological attachments. Proc R Soc A 460, 1–15.

2. Labonte, D. and Federle, W. (2015). Scaling and biomechanics of surface attachment in climbing animals. Phil Trans R Soc B 370, 20140027.

3. Rivlin, R. (1944). The effective work of adhesion. Paint Technol 9, 215–216.

4. Labonte, D., Clemente, C.J., Dittrich, A., Kuo, C.Y., Crosby, A.J., Irschick, D.J., and Federle, W. (2016). Extreme positive allometry of animal adhesive pads and the size limits of adhesion-based climbing. PNAS 113, 1297–1302.

5. Smith, J.M., Barnes, W.J.P., Downie, J.R., and Ruxton, G.D. (2006). Structural correlates of increased adhesive efficiency with adult size in the toe pads of hylid tree frogs. J Comp Physiol A 192, 1193–1204.

6. Arzt, E., Gorb, S., and Spolenak, R. (2003). From micro to nano contacts in biological attachment devices. PNAS 100, 10603–10606.

7. Bartlett, M., Croll, A., King, D., Paret, B., Irschick, D., and Crosby, A. (2012). Looking Beyond Fibrillar Features to Scale Gecko-Like Adhesion. Adv Mater 24, 1078–83.

8. Autumn, K., Dittmore, A., Santos, D., Spenko, M., and Cutkosky, M. (2006). Frictional adhesion: a new angle on gecko attachment. J Exp Biol 209, 3569–3579.

9. Endlein, T., Ji, A., Samuel, D., Yao, N., Wang, Z., Barnes, W.J.P., Federle, W., Kappl, M., and Dai, Z. (2013). Sticking like sticky tape: tree frogs use friction forces to enhance attachment on overhanging surfaces. J R Soc Interface 10, 20120838.

10. Labonte, D. and Federle, W. (2016). Biomechanics of shear-sensitive adhesion in climbing animals: peeling, pre-tension and sliding-induced changes in interface strength. J R Soc Interface 13, 20160373.

11. Federle, W. and Labonte, D. (2019). Dynamic biological adhesion: mechanisms for controlling attachment during locomotion. Phil Trans R Soc B In press.

12. Federle, W., Rohrseitz, K., and Hölldobler, B. (2000). Attachment forces of ants measured with a centrifuge: better ‘wax-runners’ have a poorer attachment to a smooth surface. J Exp Biol 203, 505–512.

13. Labonte, D. and Federle, W. (2015). Rate-dependence of ‘wet’ biological adhesives and the function of the pad secretion in insects. Soft Matter 11, 86618673.

14. Labonte, D. and Federle, W. (2013). Functionally different pads on the same foot allow control of attachment: stick insects have load-sensitive “heel” pads for friction and shear-sensitive “toe” pads for adhesion. PLoS One 8, e81943.

15. Jagota, A. and Hui, C. (2011). Adhesion, friction, and compliance of bio-mimetic and bio-inspired structured interfaces. Mat Sci Eng R 72, 253–292.

16. Autumn, K. and Hansen, W. (2006). Ultrahydrophobicity indicates a non-adhesive default state in gecko setae. J Comp Physiol 192, 1205–1212.

17. Gravish, N., Wilkinson, M., and Autumn, K. (2008). Frictional and elastic energy in gecko adhesive detachment. J R Soc Interface 5, 339–348.

18. Federle, W. and Endlein, T. (2004). Locomotion and adhesion: dynamic control of adhesive surface contact in ants. Arthropod Struct Dev 33, 67–75.

19. Bullock, J.M.R., Drechsler, P., and Federle, W. (2008). Comparison of smooth and hairy attachment pads in insects: friction, adhesion and mechanisms for direction-dependence. J Exp Biol 211, 3333–3343.

20. Clemente, C.J. and Federle, W. (2008). Pushing versus pulling: division of labour between tarsal attachement pads in cockroaches. Proc R Soc B 275, 1329–1336.

21. Tian, Y., Pesika, N., Zeng, H., Rosenberg, K., Zhao, B., McGuiggan, P., Autumn, K., and Israelachvili, J. (2006). Adhesion and friction in gecko toe attachment and detachment. PNAS 103, 19320–19325.

22. Chen, B., Wu, P., and Gao, H. (2009). Pre-tension generates strongly reversible adhesion of a spatula pad on substrate. J R Soc Interface 6, 529–537.

23. Begley, M.R., Collino, R., Israelachvili, J.N., and McMeeking, R.M. (2013). Peeling of a tape with large deformations and frictional sliding. J Mech Phys Solids 61, 1265–79.

24. Kaelble, D. (1960). Theory and analysis of peel adhesion: bond stresses and distributions. T Soc Rheol 4, 45–73.

25. Pesika, N., Tian, Y., Zhao, B., Rosenberg, K., Zeng, H., McGuiggan, P., Autumn, K., and Israelachvili, J. (2007). Peel-zone model of tape peeling based on the gecko adhesive system. J Adhesion 83, 383–401.

26. Kendall, K. (1975). Thin-film peeling – the elastic term. J Phys D: Appl Phys 8, 1449–1452.

27. Gorb, S., Gorb, E., and Kastner, V. (2001). Scale effects on the attachment pads and friction forces in syrphid flies. J Exp Biol 204, 1421–1431.

28. Dirks, J.H. and Federle, W. (2011). Mechanisms of fluid production in smooth adhesive pads of insects. J R Soc Interface 8, 952–60.

29. Betz, O. (2010). Adhesive Exocrine Glands in Insects: Morphology, Ultrastructure, and Adhesive Secretion. In Biological Adhesive Systems, J. Byern and I. Grunwald, eds. (Springer), pp. 111–152.

30. Drechsler, P. and Federle, W. (2006). Biomechanics of smooth adhesive pads in insects: influence of tarsal secretion on attachment performance. J Comp Physiol A 192, 1213–1222.

31. Hutt, W. and Persson, B. (2016). Soft matter dynamics: Accelerated fluid squeeze-out during slip. J Chem Phys 144, 124903.

32. Federle, W., Riehle, M., Curtis, A.S., and Full, R. (2002). An Integrative Study of Insect Adhesion: Mechanics and Wet Adhesion of Pretarsal Pads in Ants. Integr Comp Biol 42, 1100–1106.

33. Warton, D.I., Wright, I.J., Falster, D.S., and Westoby, M. (2006). Bivariate line-fitting methods for allometry. Biol Rev 81, 259–291.

34. Egset, C., Hansen, T., Le Rouzic, A., Bolstad, G., Rosenqvist, G., and Pélabon, C. (2012). Artificial selection on allometry: change in elevation but not slope. J Evol Biol 25, 938–948.

35. Hansen, T.F. and Bartoszek, K. (2012). Interpreting the evolutionary regression: the interplay between observational and biological errors in phylogenetic comparative studies. Syst Biol 61, 413–425.

36. Pelabon, C., Firmat, C., Bolstad, G.H., Voje, K.L., Houle, D., Cassara, J., Rouzic, A.L., and Hansen, T.F. (2014). Evolution of morphological allometry. Ann NY Acad Sci 1320, 58–75.

37. Warton, D.I., Duursma, R.A., Falster, D.S., and Taskinen, S. (2012). smatr - an R package for estimation and inference about allometric lines. Methods Ecol Evol 3, 257–259.

